# Role of Physicochemical Properties of Protein in Modulating the Nanoparticle-Bio interface

**DOI:** 10.1101/484972

**Authors:** Sunandan Dhar, Vishesh Sood, Garima Lohiya, Harini Devenderan, Dhirendra S. Katti

## Abstract

Nanoparticles, on exposure to the biological milieu, tend to interact with macromolecules to form a biomolecular corona. The biomolecular corona confers a unique biological identity to nanoparticles, and its protein composition plays a deterministic role in the biological fate of nanoparticles. The physiological behavior of proteins stems from their physicochemical aspects including surface charge, hydrophobicity, and structural stability. However, there is insufficient understanding about the role of physicochemical properties of proteins in biomolecular corona formation. We hypothesized that the physicochemical properties of proteins would influence their interaction with nanoparticles and have a deterministic effect on nanoparticle-cell interactions. To test our hypothesis, we used model proteins from different structural classes to understand the effect of secondary structure elements of proteins on the nanoparticle-protein interface. Further, we modified the surface of proteins to study the role of protein surface characteristics in governing the nanoparticle-protein interface. For this study, we used mesoporous silica nanoparticles as a model nanoparticle system. We observed that the surface charge of proteins governs the nature of the primary interaction as well as the extent of subsequent secondary interactions causing structural rearrangements of the protein. We also observed that the secondary structural contents of proteins significantly affected both the extent of secondary interactions at the nanoparticle-protein interface and the dispersion state of the nanoparticle-protein complex. Further, we also studied the interactions of different protein-coated nanoparticles with different types of cell (fibroblast, carcinoma, and macrophage). We observed that different cells internalized nanoparticle-protein complex as a function of secondary structural components of the protein. The type of model protein had a significant effect on their internalization by macrophages. Overall, we observed that the physicochemical characteristics of proteins had a significant role in modulating the nanoparticle-bio-interface at the level of both biomolecular corona formation and nanoparticle internalization by cells.

## Introduction

Nanoparticles have a unique potential for medical use since their size allows for interactions with biomacromolecules. Therefore, the primary focus of biomedical research is engineering nanoparticles to modulate their interactions with specific biomacromolecules and harness their potential for diagnostic, imaging and therapeutic applications [1, 2]. The safe and efficacious biomedical application of nanoparticles demands an in-depth understanding interface between nanoparticles and biological systems. On administration into the body, nanoparticles rapidly interact with biomacromolecules present in biological milieus such as blood plasma [3] resulting in the formation of a biomolecular corona. The biomolecular corona confers a unique biological identity to nanoparticles [4]. The physicochemical characteristics of both nanoparticles, as well as the biological components, determine the identity and quantity of biomolecules involved in the formation of biomolecular corona [5]. Although the biomolecular corona comprises of proteins, lipids, polysaccharides, and nucleic acids; proteins are the major component of this corona [6]. Proteins present in biomolecular corona significantly alters the interaction between nanoparticles and cells affecting both nanoparticle uptake and cellular behavior [7]. Protein composition of biomolecular corona has a deterministic role in the biological behaviour of nanoparticles including nanoparticle internalization by cells [8], circulation time in systemic circulation[9], and biodistribution or tissue-tropism of nanoparticles [10]. Specific plasma proteins which play a deterministic role for the biological fate of nanoparticles have also been identified [11]. The composition of biological milieus in individuals alters with physiological or pathological changes. The protein corona composition shows significant changes with conditions such as hyperglycemia, hypercholesterolemia, pregnancy and blood cancer can influence the biological response to nanoparticles [12]. Since the specific nature of changes to proteins due to these medical conditions is not well characterized, there is a lack of fundamental mechanistic understanding of how protein properties influence nanoparticle-bio interactions in such medical conditions. Thus, it is imperative to understand the fundamental aspects of nanoparticle-protein interactions to develop safe and efficacious nanotherapeutic modalities.

The protein adsorption on nanoparticle surface is modulated by both physicochemical characteristics of nanoparticles [13] and environmental factors such as the temperature of incubation [14], the time of incubation [15], and sheer-stress due to the flow of blood in vessels [16]. The protein-nanoparticle interaction is different than colloidal interactions as proteins can undergo structural rearrangements post interaction [17]. The heterogeneity in the biomolecular corona of different nanoparticles primarily arises from the heterogeneity of surface properties of proteins [18]. It is thus important to identify specific physicochemical properties of proteins which may be responsible for the differential impact of the biomolecular corona of a nanoparticle. However, since blood plasma consists of proteins with diverse physicochemical properties, it is difficult to resolve the effect of individual properties. If instead, the interactions of nanoparticles with a single model protein is analysed, we can understand the effect of the physicochemical properties of that model protein in isolation. Also, the effect of these specific properties can then be extrapolated to other proteins, thereby enabling a more generalized understanding of the process. We have employed a similar strategy in this study to investigate the role of protein physicochemical properties in modulating the nanoparticle-bio-interface.

Mesoporous silica nanoparticles (MSN) have been used in biomedical applications as a drug carrier or for diagnostic applications [19]. Therefore, we selected MSN as a model nanoparticle system for studying nanoparticle-bio interactions. We elucidated the role of protein physicochemical properties including surface charge and secondary structure content in governing MSN-protein interactions. Further, we also observed a significant effect of the nature of the nanoparticle-protein-interface in altering the interactions of protein-coated MSN with different types of cells.

## Materials and Methods

### Synthesis of Mesoporous silica nanoparticles

The detailed procedure for synthesis and characterization of mesoporous silica nanoparticles (MSN) are provided in supplementary section.

### Protein modification and characterization

Bovine serum albumin (BSA), lysozyme (Lsz) and immunoglobulin G (IgG) were procured from Sigma-Aldrich (Missouri, USA).

#### Glycation of BSA

A non-enzymatic process known as Maillard reaction was used for the glycation of BSA as described previously [20]. Briefly, 300 µM of BSA and 220 mM of dextrose were dissolved in 67 mM sodium phosphate buffer (pH 7.4) containing 0.01% (w/v) sodium azide. The solution was sterile filtered using 0.2 µm membrane filter (Whatman plc, Maidstone, UK). This solution was then incubated at 37°C for 14 days. After incubation, the solution was dialyzed against ultra-pure (type I) water (Merck Millipore, Massachusetts, USA) and stored at 4°C. Glycation was confirmed by measuring fluorescence of advanced glycation end-products (excitation at 370 nm and emission at 440 nm) [21]. The fluorescence intensity of glycated protein was compared to that of the native protein (control), which was incubated without dextrose under the same conditions.

#### Cationization of BSA

BSA was cationized by conjugating carboxyl groups of the acidic residues (aspartic acid and glutamic acid) on the surface of the protein to amine functional groups as described previously [22]. Briefly, 85 µL of hexane diamine was diluted in 3 mL of type I water and pH was adjusted to 6.5 using 1 M hydrochloric acid. This solution was then added drop-wise to a stirring solution of 150 µM BSA and 50 µM of 1-ethyl-3-(3-dimethylaminopropyl) carbodiimide (EDC) was used to catalyse the reaction. The mixture was stirred for 5 hrs at room temperature, following which EDC was once again added, and was left to stir for 6 more hrs. The pH was measured and adjusted to 6.5 at regular intervals during the reaction. Finally, the solution was dialyzed against ultra-pure (type I) water and stored at 4°C.

#### Quantification of protein concentration

Protein concentrations were determined by measuring absorbance at 280 nm using Take 3 microspot plate and a multimode spectrophotometer (Synergy H4, BioTek Instruments, Vermont, USA).

#### Surface charge of proteins

The surface charge of proteins was determined by measuring the zeta potential of the protein solutions (2 mg⋅mL^−1^). The zeta potential was calculated from the electrophoretic mobility of the proteins measured by laser Doppler velocimetry (LDV) method using a Zetasizer ZS90 (Malvern Instruments, Malvern, UK).

### Protein secondary structure content

The secondary structure of proteins was measured using a J-815 Circular Dichroism (CD) spectropolarimeter (JASCO, Oklahoma, USA) fitted with a thermostatic cell holder for temperature control. The instrument was purged with nitrogen gas before use and a constant nitrogen flow rate (5 litres min^−1^) was maintained during the experiments. The measurements were performed at a constant protein concentration (0.2 mg⋅mL^−1^) in a quartz cuvette of 1 mm pathlength at 25°C. The ellipticity was recorded between 190 to 250 nm wavelength, taking ultra-pure (type-I) water as the baseline. Data from three measurements were averaged for each sample. The ellipticity values were converted from machine units to ∆ε by normalizing to protein concentration, mean residue weight, and the pathlength of the cuvette. The secondary structure content of proteins was estimated by fitting the measured spectra to a reference dataset using CONTIN algorithm [23] on an online tool DichroWeb [24].

### Interaction between protein and nanoparticles

#### Quantification of protein adsorption

To quantify protein adsorption onto nanoparticle, the binding constants for protein-nanoparticle interaction were determined. The same volume of MSN suspension (1.0 mg⋅mL^−1^) was mixed with increasing concentrations of protein solution (0.0625 to 1.0 mg⋅mL^−1^) and incubated at 37°C for 24 hrs. The protein-bound MSN were then pelleted down by centrifugation (14000 RCF for 30 mins) and the concentration of unbound protein in the supernatant was quantified using the o-pthalaldehyde (OPA) assay with minor modifications [25]. Briefly, 50 µL of sample was mixed with 200 µL of OPA reagent (10 mg OPA, 100 µL ethanol, 5 µL β-mercaptoethanol, and 5 mL of 100 mM carbonate-bicarbonate buffer of pH 10.5) in a black opaque, non-binding 96 well-plate (Corning Inc., New York, USA). The mixture was stirred for 2 mins, following which fluorescence was measured at excitation and emission wavelengths of 340 and 455 nm respectively using multimode spectrophotometer (Synergy H4). The fluorescence intensity was correlated to protein concentration using a standard curve of the same protein. The unbound protein concentration was then subtracted from the concentration of free protein to estimate the amount of protein bound to the nanoparticles. The adsorption data was fitted to the Langmuir isotherm (equation 1) for the estimation of the binding constants for protein-nanoparticle interaction. The quantities in equation are as follow: *A* is amount of adsorbed protein (mg) per mg of MSN, *A*_*max*_ is the maximum amount of protein adsorbed at saturation (mg), [*P*] is the protein concentration (mg⋅mL^−1^), *k*_*D*_ is the dissociation constant of the protein-MSN interaction (mg⋅mL^−1^), and *n* is the Hill coefficient indicating cooperativity of binding.

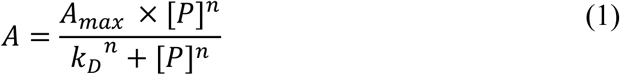

#### Temperature-dependent circular dichroism spectroscopy

The effect of nanoparticle-protein interaction on the secondary structure of the proteins was studied using CD spectroscopy [26]. Briefly, 500 µL of MSN suspension (1.0 mg⋅mL^−1^) was incubated with 500 µL of protein solution (0.2 mg⋅mL^−1^) at 37°C for 24 hrs. The concentration ratio of MSN and protein was selected so as so minimize the amount of unbound protein in the mixture, as per the values of *A*_*max*_ obtained from protein adsorption experiments. The ellipticity of proteins between 190-250 nm was recorded in the presence and absence of MSN was measured using a J-815 Circular Dichroism spectropolarimeter (JASCO, Oklahoma, USA) at temperature range of 25°C to 90°C with an interval of 5°C. Ultra-pure (type-I) water was used as the baseline for free proteins, while MSN suspension (1.0 mg⋅mL^−1^) was used as the baseline for adsorbed proteins. The CD data was then used to calculate thermodynamics of unfolding of the proteins. First, the fraction of protein unfolded at a given temperature was calculated using equation 2 where *K*_*unf*_ is the fraction of protein unfolded, [*U*] is the concentration of unfolded protein (mg⋅mL^−1^), [*F*] is the concentration of folded protein (mg⋅mL^−1^), *θ*_*T*_ is the ellipticity of protein at a given temperature (Δ*ε*), *θ*_*F*_ is the ellipticity of the fully folded protein in the native state (Δ*ε*), *θ*_*U*_ is the ellipticity of protein in the fully unfolded state (Δ*ε*). Ellipticity values at 222 nm for BSA and Lsz, and at 216 nm for IgG were recorded due to the presence of characteristic peaks at these wavelengths. For the calculations, it was assumed that proteins are in their fully-folded, native state at 25°C, and in the fully-unfolded state at 90°C. The enthalpy and entropy of protein unfolding in presence and absence of MSN were estimated using the van’t Hoff equation (equation 3). Finally, the melting temperature is the temperature at which the protein is half-folded and thus free energy of protein unfolding is equal to zero. This value was calculated according to equation 4. The quantities in the equations are as follows: ∆*G*_*unf*_ is the free energy of protein unfolding (J⋅mol^−1^), *R* is the gas constant (8.314 J⋅mol^−1^⋅K^−1^), *T* is the temperature (K), Δ*H* is the enthalpy required for unfolding (J⋅mol^−1^), Δ*S* is the entropy required for unfolding (J⋅mol^−1^⋅K^−1^), and *T*_*m*_ is the protein melting temperature (K), at which the protein is half unfolded.

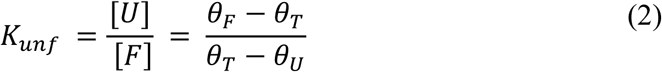

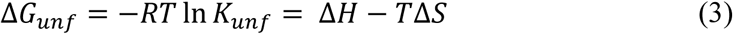

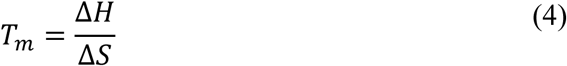

Changes in secondary structure of proteins with temperature in presence and absence of MSN was also estimated by fitting the measured spectra to reference dataset using DichroWeb [24].

### Interaction between cells and protein-coated-nanoparticles

#### Cell culture

Three cell lines: NIH/3T3 (mouse fibroblast), Ehrlich-Lettre ascites strain E (EAC-E) (mouse carcinoma) and RAW264.7 (mouse macrophage) were obtained from National Centre for Cell Science (NCCS), Pune. Cells were cultured in a 1:1 mixture of Ham’s F12 nutrient mixture and Dulbecco’s modified Eagle media (DMEM) supplemented with 10% (v/v) FBS and antimicrobials (100 U⋅mL^−1^ penicillin-streptomycin, 2.5 µg⋅mL^−1^ amphotericin B and 2.5 µg⋅mL^−1^ ciprofloxacin). Cells were incubated in a humidified atmosphere containing 5% CO_2_ at 37°C. NIH/3T3 and EAC-E cells were detached from T-25 flasks using 0.05% trypsin-EDTA in DMEM, while RAW264.7 cells were detached by incubating with 10 mM EDTA in 1X PBS at 4°C for 10 mins followed by scraping. Cell count and viability were determined using Muse® Cell Analyzer (EMD Millipore, Massachusetts, USA). Cells were seeded on multi-well plates (Corning, New York, USA) at a density of 25,000 cells⋅cm^−2^ for all experiments.

#### Cellular internalization of nanoparticles

Internalization of MSN into cells was measured using flow cytometry. TRITC-tagged MSN were suspended in 2X serum-free media, mixed with an equal volume of aqueous protein solution, and allowed to equilibrate for 2 hrs. This protein-coated MSN suspension was then added to cells seeded on 24-well plates and incubated at 37°C. At the required time point, MSN suspension was aspirated, and cells were washed three times using 1X PBS. Cells were then detached using trypsin-EDTA for NIH/3T3 and EAC-E, and for RAW264.7 cells were scraped to obtain a cell suspension. The cell suspension was pelleted down, washed using 1X PBS and re-suspended in PBS. Muse® Cell Analyzer (MilliporeSigma, USA) was used to detect TRITC fluorescence in cells to quantify internalized MSN.

#### Determination of surface charge density of the cell membrane

The surface charge density of the cell membrane before and after interaction with protein-coated MSN were determined as described previously [27]. MSN were suspended in 2X serum-free media and mixed with an equal volume of protein solution, and allowed to equilibrate at 37°C for 2 hrs. This protein-coated MSN suspension was then added to cells that were seeded in 6-well plates. After 6 hrs, the MSN suspension was aspirated, and cells were washed three times with 1X PBS. Cells were then trypsinized or scraped and resuspended in an isotonic solution (240 mM sucrose, 20 mM glucose, and 20 mM sodium chloride). The electrophoretic mobility of the suspended cells under an applied electrical field was measured using a Zetasizer ZS90. The surface charge density was calculated from electrophoretic mobility using equations 5 and 6 where *σ* is surface charge density (C⋅m^−2^), *η* is the viscosity of the medium (kg⋅m^−1^⋅s^−1^), *μ* is the electrophoretic mobility of the cells (m^2^⋅V^−1^⋅s^−1^), *d* is the thickness of the diffuse layer (m), *ε* ∙ *ε*_0_ is the permittivity of the medium (C⋅V^−1^⋅m^−1^), *R* is the gas constant (8.314 J⋅mol^−1^⋅K^−1^), *T* is the temperature of the medium (310 K), *I* is the ionic strength of the medium (20 mM), *F* is the Faraday constant (96485.33 C⋅mol^−1^).

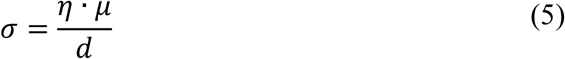

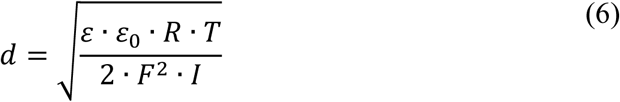

### Statistical analysis

Three experimental replicates were performed for all experiments and mean values are represented in all graphs and tables, with error bars representing the standard error of the mean. A confidence interval of 95% was used to evaluate statistical significance. Statistical analysis of experimental data was performed using the GraphPad Prism software, using either the Student’s t-test or analysis of variance (ANOVA) with the Tukey post-hoc test. Three-way ANOVA was performed using the IBM SPSS Statistics software.

## Results and Discussion

### Selection and characterization of model proteins

Physicochemical properties of proteins have a deterministic effect on their interaction at solid-liquid interfaces [28, 29]. Surface charge is one of the critical physicochemical properties of proteins which influences the extent and kinetics of protein adsorption onto planar surfaces [30] and is a critical determinant for nanoparticle-bio interfaces [31, 32]. The surface of the protein, especially, ligand binding sites, is derived from the structure of the proteins [33]. However, the role of protein structure in modulating the nanoparticle-bio interface is not well understood. The secondary structure components of proteins form the basis for Structural Classification of Proteins (SCOP) [34]. To understand the correlation between protein structure and nanoparticle-protein interaction, we selected model proteins from different SCOP classes: i) bovine serum albumin (BSA) as an all α-helix protein, ii) immunoglobulin G (IgG) as an all β-sheet protein, and iii) lysozyme (Lsz) as an α-helix and β-sheet protein.

Further, we studied the effect of glycation of the protein surface. Glycation of proteins commonly occurs in hyperglycaemic individuals, and thus its influence on nanoparticle interactions can have an impact in the biomedical applications of nanoparticles [35]. BSA was incubated with excess glucose, following which a significant increase in fluorescence intensity of glycated BSA (gly-BSA) at 440 nm when excited at 370 nm was observed (Figure 1A) indicating the formation of advanced glycation end-products [20].

**Figure 1:**
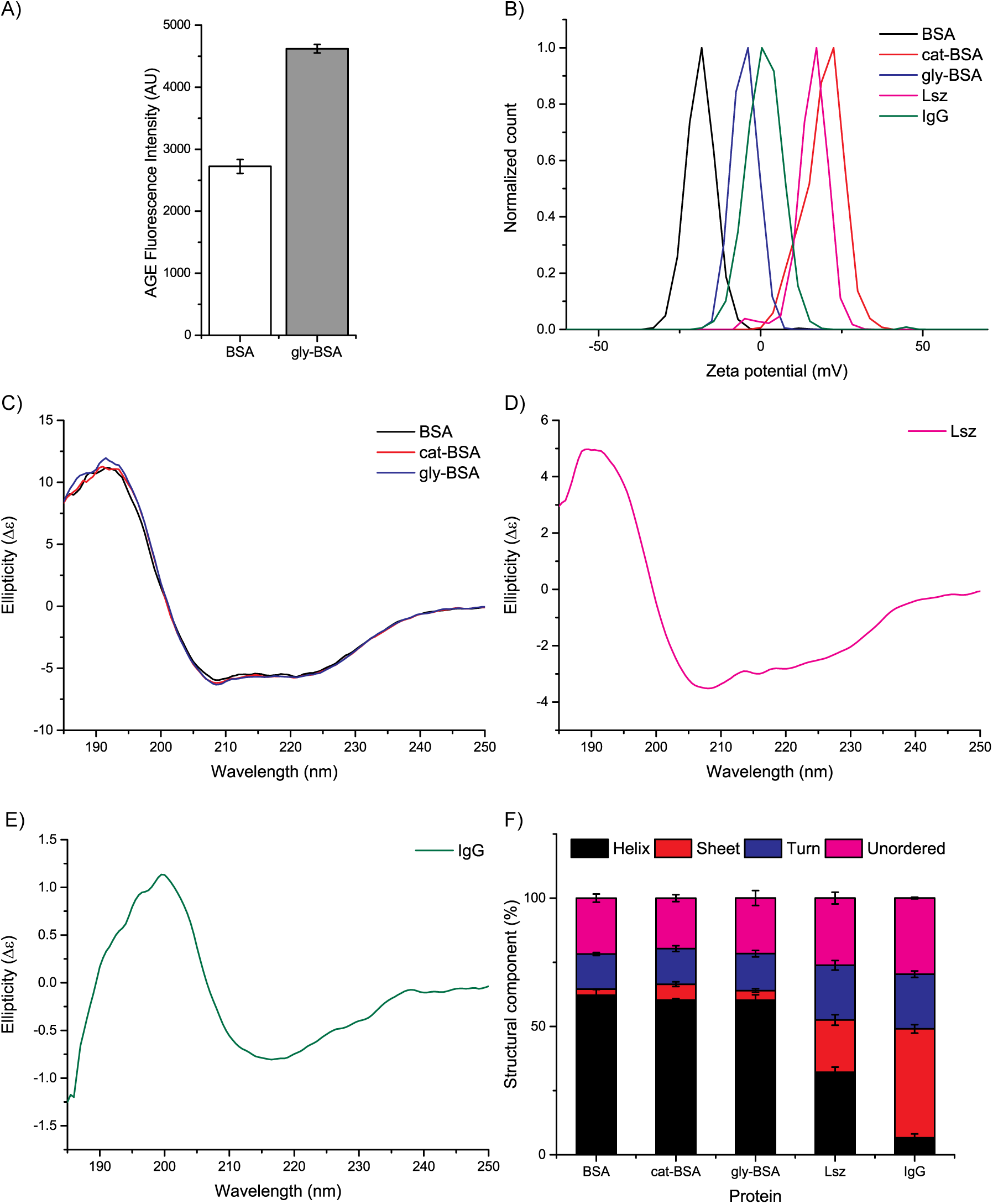
Physicochemical characterization of model proteins. A) Fluorescence of advanced glycation end-products (AGEs) of native and glycated BSA. B) Zeta potential distributions of the model proteins. C-E) Circular dichroism spectra of C) native BSA, cationized BSA (cat-BSA), and glycated BSA (gly-BSA); D) lysozyme (Lsz); and E) Immunoglobulin (IgG). F) Percentage content of secondary structural components of the model proteins.

The above model proteins differ both in surface charge and secondary structure. In order to make a more direct comparison and resolve the specific effect of charge, we modified the surface charge of BSA by conjugating cationic amine groups to surface carboxylic acids of BSA. We confirmed the altered surface charge by measuring the zeta potential of the proteins in aqueous solution (Figure 1B). Native BSA had a zeta-potential of −18.6 mV, which increased to +19.3 mV for cationized BSA (cat-BSA). We also measured the zeta potential of other model proteins. The zeta potential of gly-BSA, Lsz, and IgG was −4.9 mV, +13.4 mV, and +1.1 mV respectively. Thus, BSA and gly-BSA had an anionic surface charge, while cat-BSA and Lsz had a cationic surface charge, and IgG had a nearly neutral surface charge.

We also studied the secondary structure content of the model proteins using Circular dichroism (CD) spectroscopy (Figure 1C-E). The CD spectra for BSA, gly-BSA, and cat-BSA were overlapping (Figure 1C). We observed no significant variations in the secondary structure of BSA due to the modifications (Supplementary Table S1). These proteins had a high content of α-helices and a low content of β-sheets (Figure 1F). Lsz had a lower content of helices and a higher content of sheets. IgG had a significantly high content of sheets and low content of helices. There was a significant difference in the α-helix and β-sheet content of BSA (native and modified), Lsz and IgG (Supplementary Table S1). These five model proteins were used to study interactions with nanoparticles.

### Interactions between nanoparticles and proteins

We used Mesoporous silica nanoparticles (MSN) synthesized using the reverse-microemulsion technique as a model nanoparticle system. The synthesized MSN were monodispersed (Z average hydrodynamic diameter of 90.16 nm with PDI of 0.066) with an anionic surface charge (-42.2 mV) and formed a stable suspension in water, PBS, and cell culture media (Supplementary Figure 1).

On exposure to protein solutions, we observed an increase in the size of MSN (Supplementary Table 2). We also observed agglomeration of MSN after exposure to cat-BSA and IgG. The protein-coated MSN also showed a change in the zeta potential (Supplementary Table 2) with adsorption of BSA and gly-BSA causing a decrease in zeta potential, whereas, adsorption of cat-BSA, Lsz, and IgG causing a change in the polarity of zeta potential.

To study the kinetics of protein interaction with MSN, we quantified the amount of adsorbed proteins and fitting the data to an adsorption isotherm (Figure 2). We observed significant differences between the adsorption behavior of different model proteins, indicating the influence of physicochemical properties in governing protein-nanoparticle interactions. Adsorption isotherms yielded the amount of protein bound normalized to per unit mass of nanoparticles (*A*_*max*_), the propensity of proteins to dissociate from the nanoparticle at equilibrium estimated as dissociation constant (*k*_*D*_), and binding cooperativity of proteins estimated as *n* (Figure 2F). We observed significant changes in the amount of proteins adsorbed (*A*_*max*_) for all the model proteins (Supplementary Table 2). *A*_*max*_ was significantly higher for the cationic proteins: cat-BSA (Figure 2B) and Lsz (Figure 2D) than for the anionic proteins: BSA (Figure 2A) and gly-BSA (Figure 2C). These results indicate that electrostatic interactions between proteins and the anionic MSN was a significant factor determining the amount of protein bound to nanoparticles. Our results also agree with existing literature about planar surfaces which show that favourable electrostatic interactions enhance the amount of protein adsorbed [28, 36]. We observed the highest *A*_*max*_ value for IgG (Figure 2E), which had a neutral surface charge. The adsorption of proteins at isoelectric point reaches a maximum value for both BSA [37] and IgG [38] and it is postulated to be due to loss of long-range electrostatic screening leading to increased lateral interactions between protein molecules [38]. In the case of nanoparticles, we envision that increased lateral interaction will result in increased multiple interactions between nanoparticles and proteins resulting in the formation of protein enmeshed nanoparticle agglomerates as observed for IgG-MSN (Supplementary Table 2).

**Figure 2.**
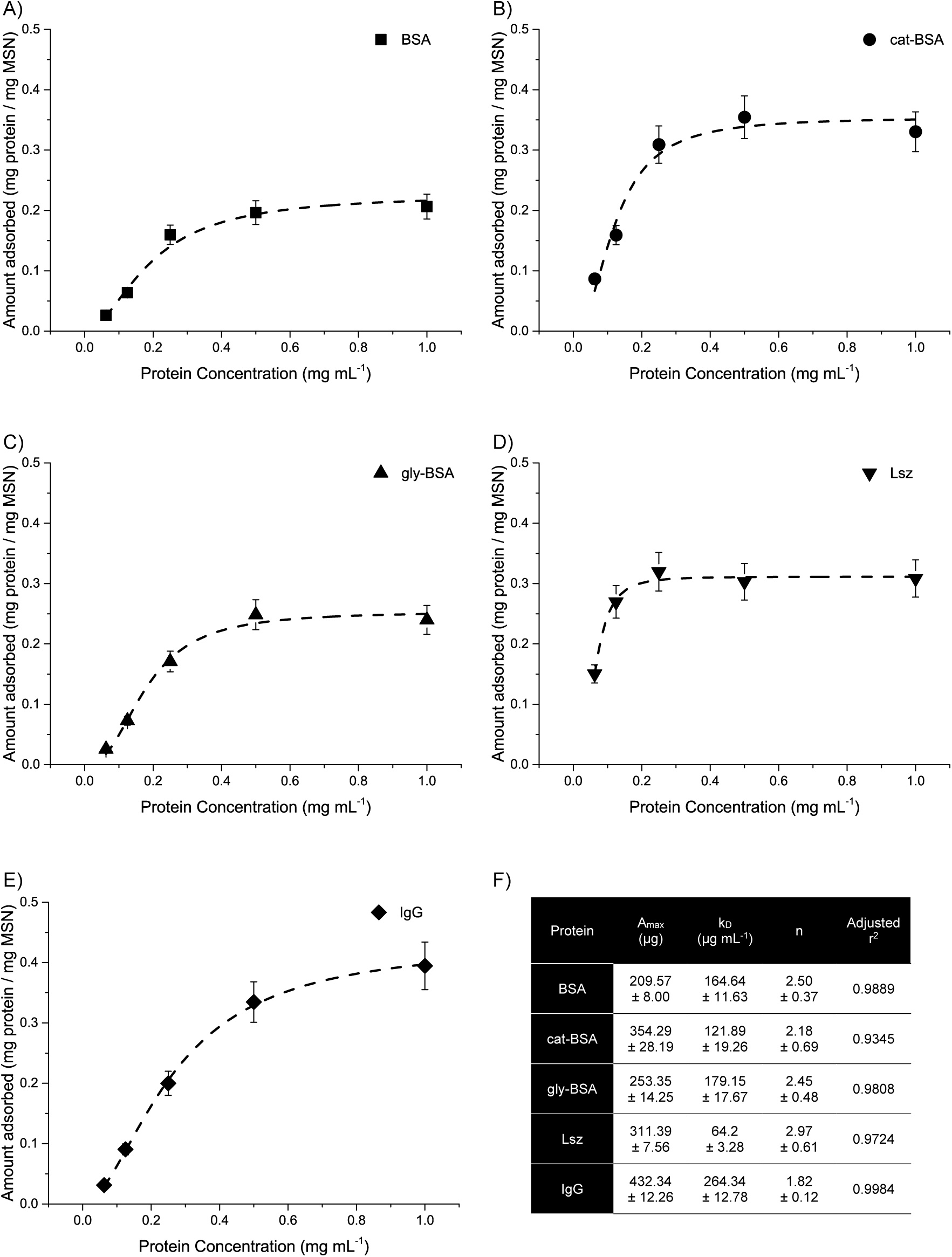
Protein adsorption isotherms. Adsorption of A) BSA, B) cat-BSA, C) gly-BSA, D) Lsz, and E) IgG onto MSN. The dashed lines represent the best fit of the data to the Langmuir adsorption isotherm. F) Adsorption parameters obtained from isotherms: maximum amount of protein adsorbed (***A***_***max***_), dissociation constant (***k***_***D***_) and binding cooperativity (***n***).

The anionic proteins (BSA and gly-BSA) had significantly higher *k*_*D*_ values, and thus lower binding affinities, than the cationic proteins (Lsz and cat-BSA). Similar results have been reported previously whereby cationization of protein surface increased the binding affinity between human serum albumin and anionic quantum [39]. We, therefore, inferred that favorable electrostatic interactions play a deterministic role in the kinetics of nanoparticle-protein interactions. Interestingly, cat-BSA had a significantly lower binding affinity than Lsz (Supplementary Table 3) despite having a similar cationic surface charge (Supplementary Figure 1) indicating that surface charge alone was not enough to explain the protein-nanoparticle interaction. Moreover, the lack of any significant differences in binding affinities of BSA and gly-BSA despite significantly more *A*_*max*_ for gly-BSA (Supplementary Table 3) indicate the importance of glycation in modulating the biomolecular corona of nanoparticles. Finally, IgG had the lowest binding affinity, despite having the highest *A*_*max*_ value due to agglomeration (Supplementary Table 2). All model proteins had positive binding cooperativity with the MSN with no significant differences in values of *n* (Figure 2F).

### Structural rearrangements in proteins after protein-nanoparticle interaction

After the initial binding of a protein to the nanoparticle surface, the protein molecules may undergo structural rearrangements in order to minimize the energy at the interface. For a macroscopic planar surface, proteins with lower structural stability undergo a greater extent of structural rearrangements after adsorption [29]. However, no existing reports study the role of protein properties on such effects during adsorption onto nanoscale, curved surfaces such as nanoparticles. The extent of these structural changes can determine the extent of desorption that the adsorbed proteins can undergo, affecting the dynamics at the nanoparticle-protein interface [40]. Structural rearrangements may also lead to the exposure of previously buried protein domains, altering the function and interactions of the protein [41]. Thus, we studied structural rearrangements of proteins post adsorption on to nanoparticles to decipher the importance of protein physicochemical properties on nanoparticle-protein interactions.

We used Circular dichroism (CD) spectroscopy to estimate changes in secondary structures of proteins after binding to MSN. We recorded the CD spectra as a function of temperature for an in-depth analysis of the structural stability of the protein in the presence and absence of MSN (Figure 3). Interestingly, α-helices of all proteins except gly-BSA were significantly affected due to interaction with MSN (Table 1). However, the contribution of MSN interaction had a small but significant effect on the secondary structure for BSA, gly-BSA, and IgG as the increase in temperature sufficiently explained changes in the secondary structure of the proteins (Table 1). For cationic proteins, cat-BSA and Lsz, the secondary structures were significantly altered due to MSN interaction as well as temperature. MSN interaction contributed significantly to the decrease in α-helices and an increase in β-sheets content of cat-BSA, as well as to an increase in α-helices and the decrease in β-sheets of Lsz. Cat-BSA and Lsz also had the highest binding affinities with MSN (Figure 2). Therefore, favorable electrostatic interactions at the nanoparticle-protein interface not only facilitated adsorption but also correlated to the extent of structural rearrangements in proteins after adsorption.

**Table 1.**
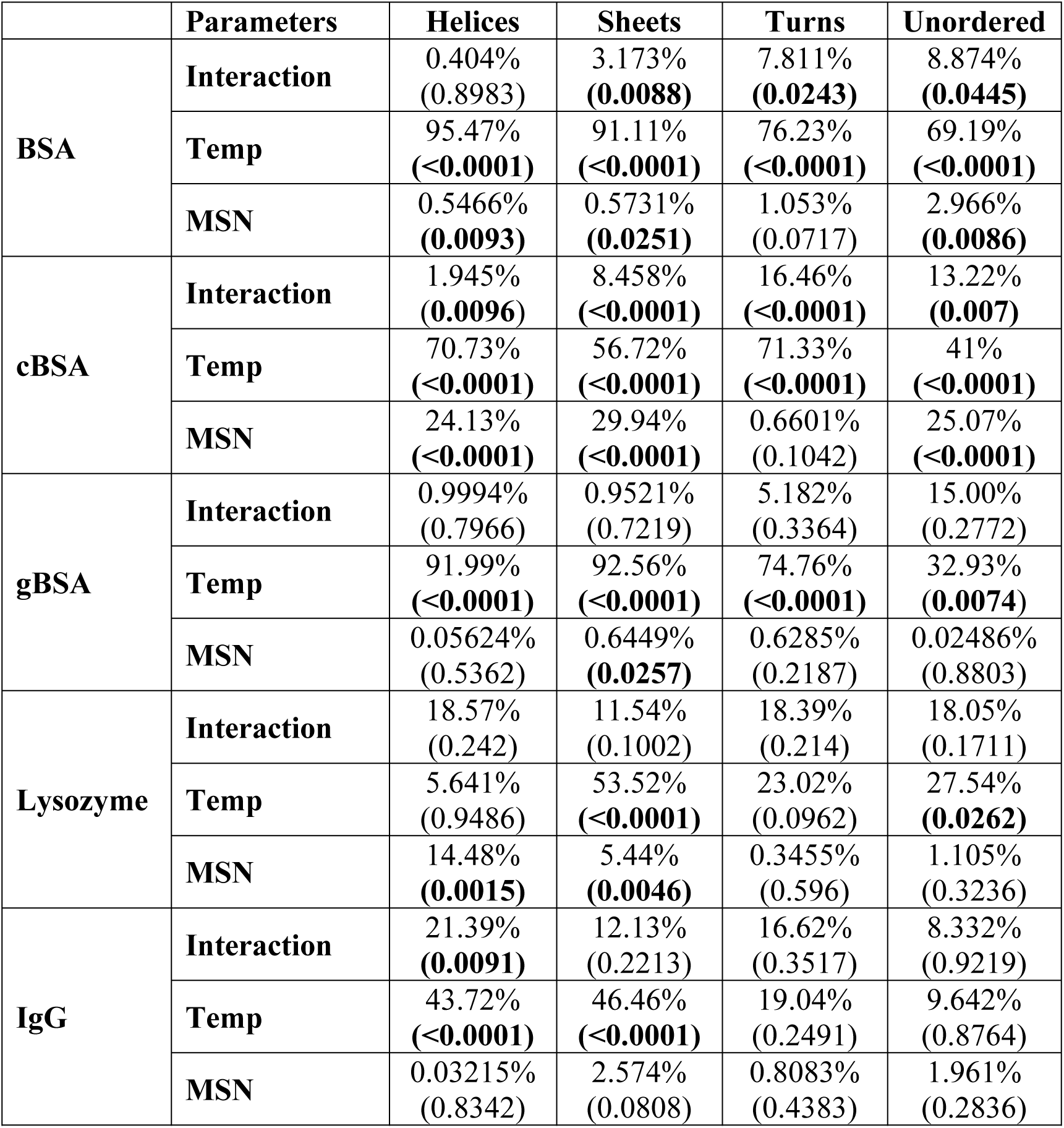
A two-way ANOVA for changes in the secondary structures of proteins with increase in temperature in presence and absence of MSN. The percentage variation explained by Temperature, presence of MSN or their interaction is provided with p-values given in parenthesis. Significant p-values are provided in bold-face.

**Figure 3.**
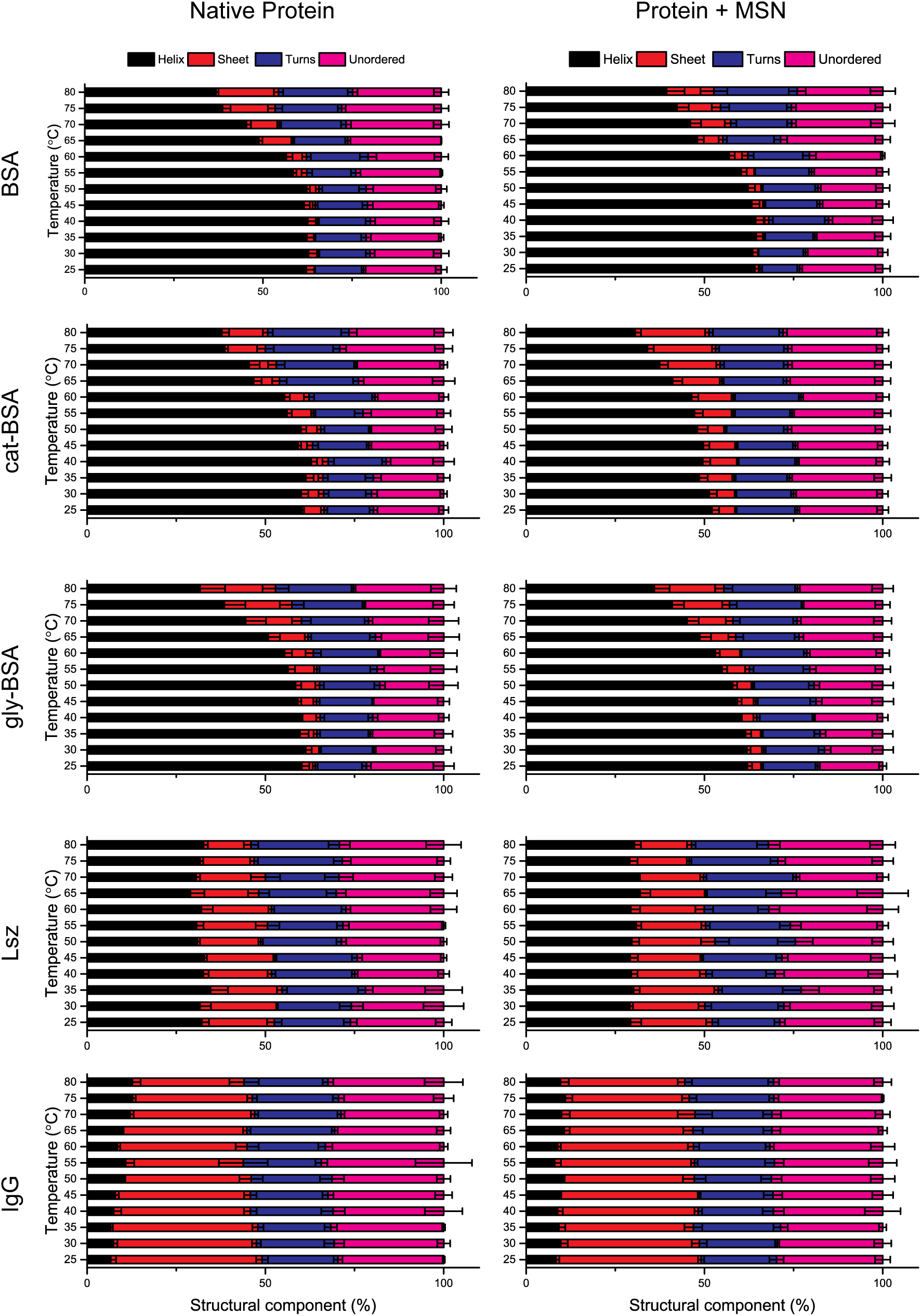
Temperature-dependent secondary structure content of native and MSN-bound proteins. Percentage content of helices, sheets, turns and unordered regions estimated from the CD spectra of native and MSN-bound proteins at increasing temperature.

The rigidity of proteins is directly related to the content of secondary structural elements, and proteins with a higher content of sheets are more resistant to structural rearrangements post adsorption [42]. We studied conformational changes in proteins after MSN interaction using fluorescence quenching (Supplementary Figure S2) [43]. IgG, with the highest sheet content, was observed to have the lowest quenching constant (*K*_*SV*_), indicating that it underwent the lowest extent of conformational change after interaction with MSN. Both BSA and Lsz had significantly lower *K*_*SV*_ values when compared to cat-BSA and gly-BSA (Supplementary Table 4). Lsz is inherently stable due to the presence of β-sheets, whereas, BSA has an anionic surface and thus had a lower binding affinity to the anionic MSN (Supplementary Table 2). Both modifications of BSA resulted in significantly higher structural deformation (Supplementary Table 4). These results indicate that glycation of proteins can significantly alter the behavior of protein during protein corona formation. While for cat-BSA, favorable electrostatic interactions with MSN resulted in a greater extent of conformational changes. Moreover, a significant structural rearrangement in cat-BSA post adsorption can explain agglomeration of cat-BSA-coated MSN (Supplementary Table S2), in contrast to LSZ-coated MSN which were colloidally stable.

We also studied the effect of the presence of MSN on the nature of protein unfolding by studying ellipticity of proteins at 201 and 208 nm [44]. The difference in the ellipticity of unfolded proteins (at 80 °C) and folded proteins (at 25 °C) at 201 nm signifies induction of β structures (sheets and turns) or unordered structures (Figure 4). The ratio of the ellipticity of unfolded proteins (at 80 °C) and folded proteins (at 25 °C) at 208 nm signifies the residual α helical content of the proteins (Figure 4). At 201 nm, a negative change in 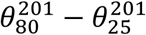 signifies induction of unordered structures, whereas, a positive change signifies induction of β structures (sheets and turns). We observed that presence of MSN significantly affected the folding behavior of BSA and gly-BSA with denaturation in the presence of MSN resulting in induction of unordered structures, whereas, for native proteins, there was an increase in β structures (Figure 4A, B). We also observed a decrease in the induction of unordered structure of IgG in MSN-bound form which can be attributed to agglomeration of MSN and IgG. We observed no significant change in the denaturation of cat-BSA and Lys. At 208 nm, the ratio 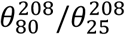 signifies changes in the α helical content with a value less than 1 indicating loss of helices and a value more than 1 indicating induction of helices. We observed that for BSA and gly-BSA, the residual helical content increased in the presence of MSN indicated by a value closer to 1 when compared to native proteins (Figure 4C, D). For cat-BSA, the residual helical content decreased significantly in the presence of MSN indicating structural rearrangements. We also observed significant changes in the denaturation behavior of IgG. Denaturation induced α helices for native IgG, whereas, denaturation resulted in a decrease of helical content in the presence of MSN primarily due to agglomeration. Interestingly, we observed no significant changes for Lsz indicating that structurally stable proteins undergo minimal structural rearrangements after adsorption on nanoparticles even when electrostatic interactions are favorable.

**Figure 4.**
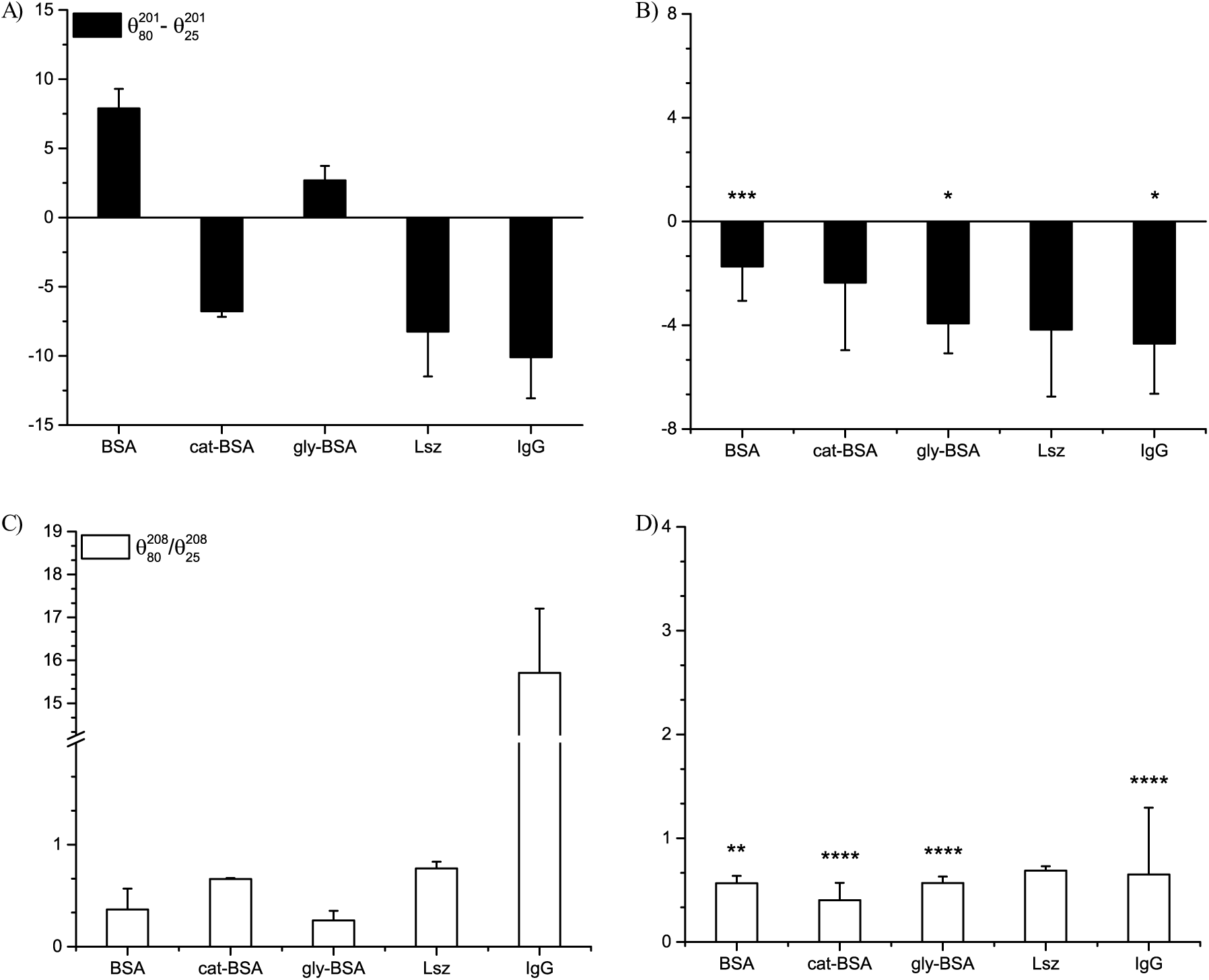
Nature of secondary structure changes in model proteins in native (A, C) and MSN bound state (B, D). 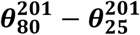 is change in the ellipticity of proteins at 201 nm in unfolded (80 °C) and folded state (25 °C). The unfolding results in induction of β structures (sheets and turns) when the value of 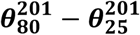 is positive while there is induction of unordered structures when the value is negative. 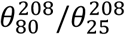 is ratio of ellipticity of proteins at 208 nm in unfolded (80 °C) and folded (25 °C). The unfolding causes an increase in the α helical content of proteins if the value of 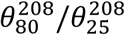 is more than 1 while α helical content decreases if the value of 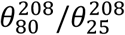 is less than 1. The value of 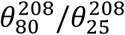 also represents residual amount of α helical content of proteins after thermal denaturation. A 2-way ANOVA analysis between presence and absence of MSN and secondary structure changes at different temperatures was used to infer the statistical significance of the results obtained. The p-values reported are * p<0.05, ** p<0.01, *** p<0.001, and **** p<0.0001.

Changes in the secondary structures may alter the thermal stability of the protein. However, there are no reports of observing such an effect for nanoparticle-protein interactions. We studied the thermal stability of proteins in the presence and absence of nanoparticles to study the contribution of nanoparticle-protein interactions in structural rearrangement of proteins. We used the rate of thermal unfolding of native and MSN-bound proteins to calculate the free energy of melting (Supplementary Figure 3), which was then used to estimate the enthalpy and entropy of unfolding (Table 2). We observed a significant loss in the thermal stability of cat-BSA and Lsz after interaction with MSN, indicated by a decrease in enthalpy and entropy of unfolding as well as lower melting temperatures. Thus, a greater extent of protein adsorption not only correlates to changes in protein secondary structure, but also to a loss in structural stability of the MSN-bound proteins. Lsz is known to be inherently more structurally stable than BSA and IgG [29], which is also reflected by the values for native proteins in Table 2. Our observations indicate that after interaction with nanoparticles, the inherent stability of Lsz is lost mainly due to the decrease in α-helical content (Table 1). However, due to a higher β-sheet content of Lsz, the structural rearrangements are not as drastic as cat-BSA. Therefore, we did not observe any agglomeration of Lsz-coated MSN (Supplementary Table 2).

**Table 2.**
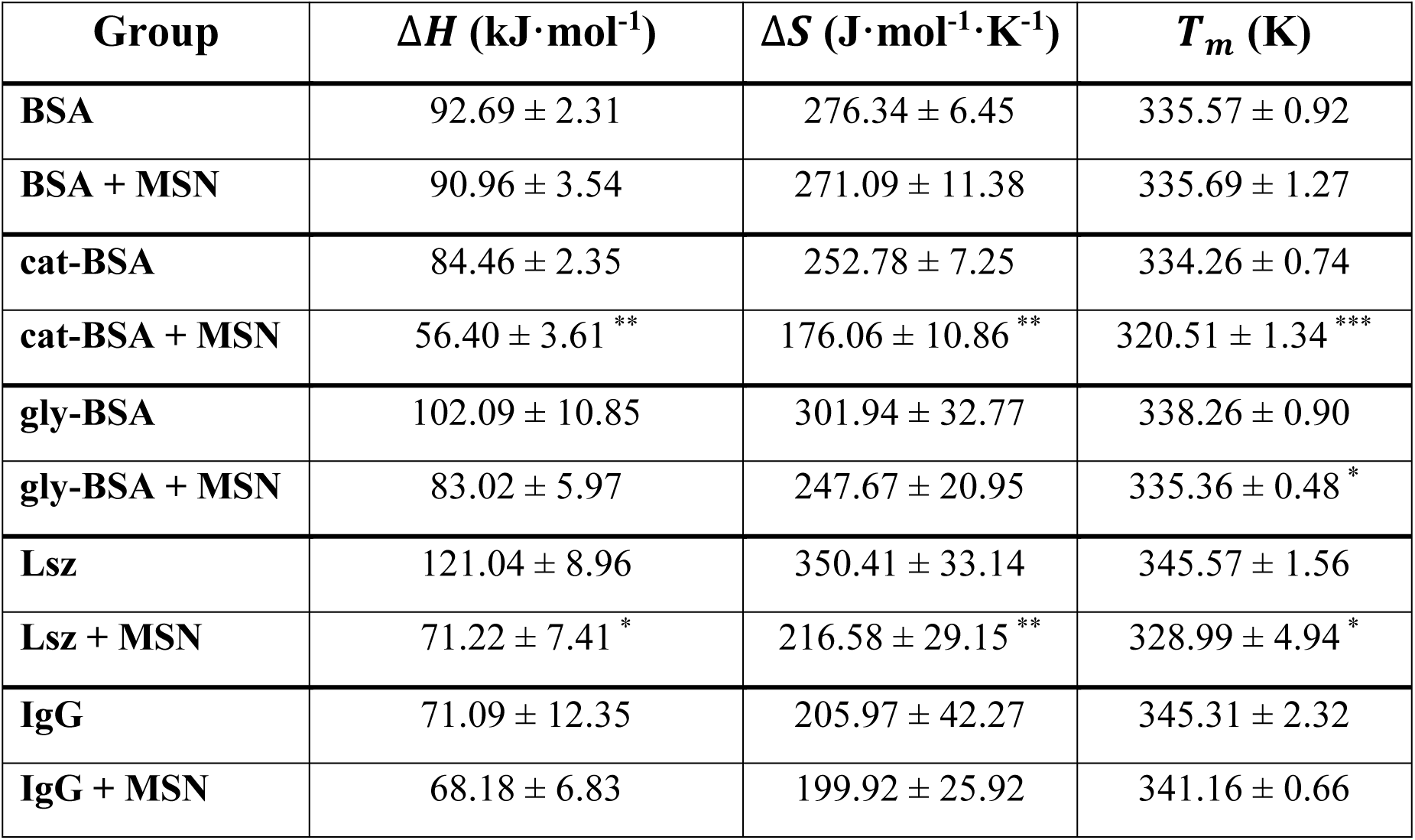
Thermal stability of native and MSN-bound proteins. Enthalpy (∆***H***) and entropy (∆***S***) of heat-induced unfolding at melting temperature (***T***_***m***_) of native and MSN-bound proteins. * indicates statistically significant difference with respect to the corresponding protein in native state (* p<0.05, ** p<0.01, *** p<0.001).

| Group | $\Delta H$ (kJ·mol<sup>-1</sup>) | $\Delta S$ (J·mol<sup>-1</sup>·K<sup>-1</sup>) | $T_m$ (K) |
| --- | --- | --- | --- |
| **BSA** | 92.69 ± 2.31 | 276.34 ± 6.45 | 335.57 ± 0.92 |
| **BSA + MSN** | 90.96 ± 3.54 | 271.09 ± 11.38 | 335.69 ± 1.27 |
| **cat-BSA** | 84.46 ± 2.35 | 252.78 ± 7.25 | 334.26 ± 0.74 |
| **cat-BSA + MSN** | 56.40 ± 3.61 ** | 176.06 ± 10.86 ** | 320.51 ± 1.34 *** |
| **gly-BSA** | 102.09 ± 10.85 | 301.94 ± 32.77 | 338.26 ± 0.90 |
| **gly-BSA + MSN** | 83.02 ± 5.97 | 247.67 ± 20.95 | 335.36 ± 0.48 * |
| **Lsz** | 121.04 ± 8.96 | 350.41 ± 33.14 | 345.57 ± 1.56 |
| **Lsz + MSN** | 71.22 ± 7.41 * | 216.58 ± 29.15 ** | 328.99 ± 4.94 * |
| **IgG** | 71.09 ± 12.35 | 205.97 ± 42.27 | 345.31 ± 2.32 |
| **IgG + MSN** | 68.18 ± 6.83 | 199.92 ± 25.92 | 341.16 ± 0.66 |

Overall, an interplay between electrostatic interactions and secondary structure of proteins play a deterministic role in defining the nanoparticle-protein interface. Glycation of proteins influences nanoparticle interaction and subsequent structural rearrangements of proteins. As we were unable to modify the surface of both Lsz and IgG by glycation, this limited the scope of the manuscript in deciphering the influence of the secondary structure of glycated proteins on nanoparticle-protein interactions.

### Interactions between protein-coated nanoparticles and cells

The nature of the protein corona around a nanoparticle influences its interactions with cells. Previous studies have explored the impact of the presence or absence of the protein corona on cellular interactions [45], as well as the effect of particular plasma proteins which can influence the internalization of nanoparticles by different types of cells [46]. However, the role of specific physicochemical properties of proteins which cause differential recognition of nanoparticles by different cell types is not well understood. To address this, we studied the role of protein secondary structure and surface properties on interactions between protein-coated nanoparticles and cells.

Keeping in mind the pluralistic nature of the cellular response to nanoparticles based on the type of cell involved, we used three types of cells to gain an in-depth understanding of interactions between protein-coated nanoparticles and cells: fibroblasts (NIH/3T3), carcinoma cells (EAC-E) and macrophages (RAW264.7). Fibroblasts are normal cell found in various tissues. Cancer cells such as carcinoma have an altered phenotype compared to healthy cells, and thus often respond differently to nanoparticles [47]. Macrophages are a type of immune cell which serves as a primary biological response to nanoparticles introduced into the bloodstream, affecting the clearance rate and biodistribution of nanoparticles [48]. All three cell lines used were from mouse origin, to avoid any species-related differences. The viability of these three cell types was not significantly affected by exposure to MSN (50 μg mL^−1^) coated with the five proteins (Supplementary Figure S4).

The cellular internalization of TRITC-tagged MSN was detected using flow cytometry (Supplementary Figure S6). We observed influence of both cell-type and protein corona on cellular internalization of nanoparticles with variations in both amount and kinetics of internalization (Figure 5). In general, we observed that the presence of a protein corona impeded the kinetics of internalization as MSN uptake saturated at a lower time point in comparison to protein-coated MSN groups (Figure 5). Our results are as per the reported literature as the presence of protein corona significantly reduces uptake of silica nanoparticles by cells [7]. A 3-way ANOVA analysis revealed that exposure time did not have a significant contribution to the variation in the data (Supplementary Table S6). On the other hand, cell type, type of protein corona, and the interaction of these two sources together contributed more than 78% of the variation in the data. Therefore, we fixed the exposure time at 6 hours for further analysis since most groups had reached saturation by this time point (Figure 6A-C). The presence of any protein corona significantly decreased the internalized amount for all cell types. Moreover, the properties of proteins significantly altered the internalization of protein-coated MSN. MSN coated with cat-BSA were internalized in the lowest quantity for all types of cells. Agglomeration of cat-BSA coated MSN (Supplementary Table 2) may lead to lower internalization due to the larger size of the complex. We observed a significant differential effect of protein physicochemical properties on the uptake of nanoparticles by macrophages (Figure 6C). Macrophages internalized Lsz-coated MSN in significantly higher amount than the other protein-coated MSN. Lsz-coated MSN was the only complex which did not agglomerate and had a cationic surface charge (Supplementary Table 2) and may thus adhere favorably to the anionic cell membrane, aiding in its cellular internalization. We observed a similar trend for carcinoma cells as well, but the variance in the data was more (Figure 6B). Also, the internalization of gly-BSA-coated MSN was higher than that of native BSA-coated MSN for macrophages, indicating that glycation of proteins may significantly influence the nanoparticles uptake by reticuloendothelial cells. As diabetes causes similar glycation of serum proteins, therefore, it might be a contraindication for nanotherapeutic interventions and needs further investigation. We speculate that increased uptake of gly-BSA-coated MSN may be due to the specific recognition of glycated proteins by the receptors for advanced glycation end-products (RAGEs) present on the surface of macrophages [35].

**Figure 5.**
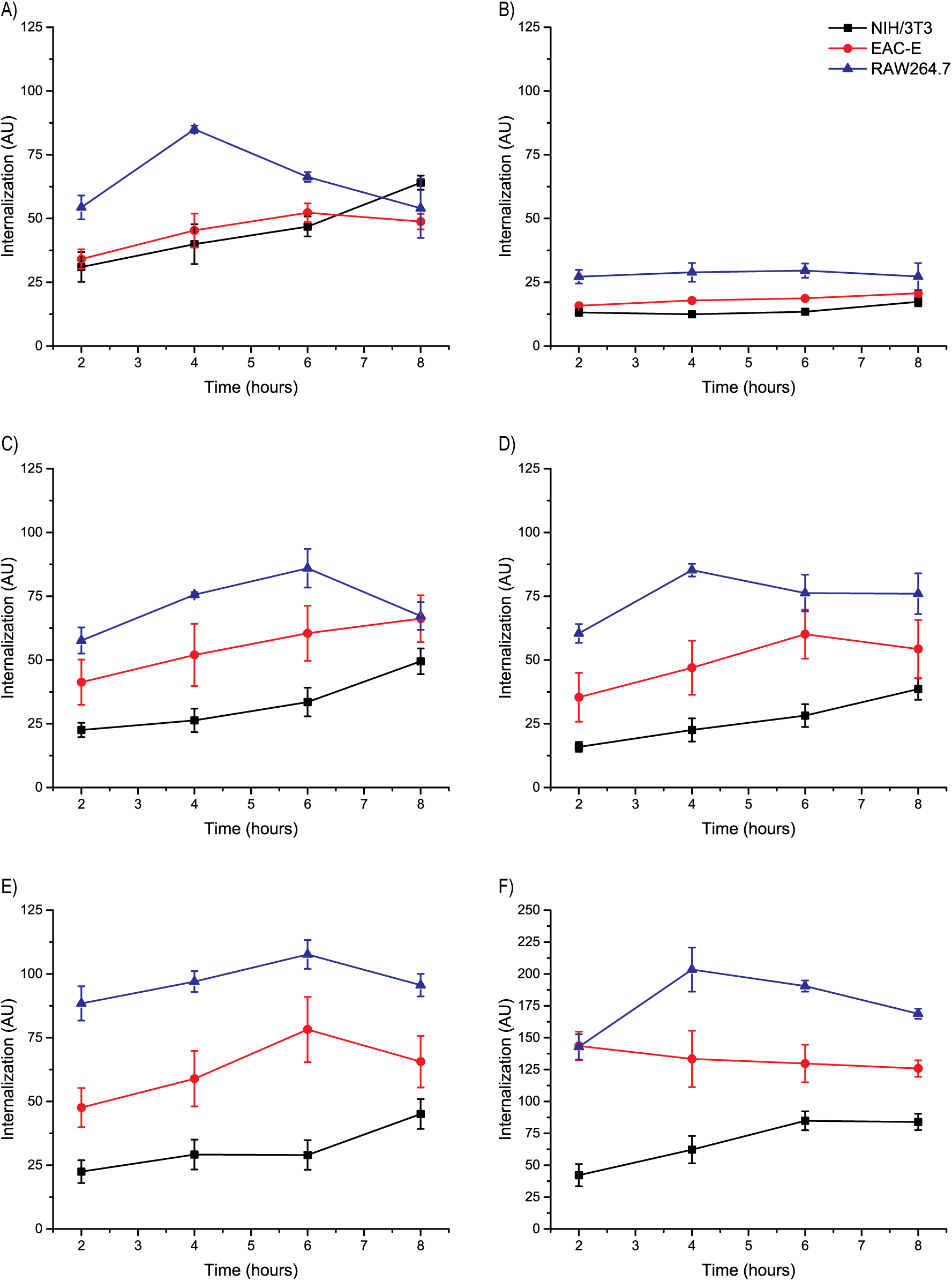
Cellular internalization kinetics. Internalization of MSN coated with A) BSA, B) cat-BSA, C) gly-BSA, D) Lsz, E) IgG, and F) bare MSN after 2, 4, 6 and 8 hrs of exposure to cells.

**Figure 6.**
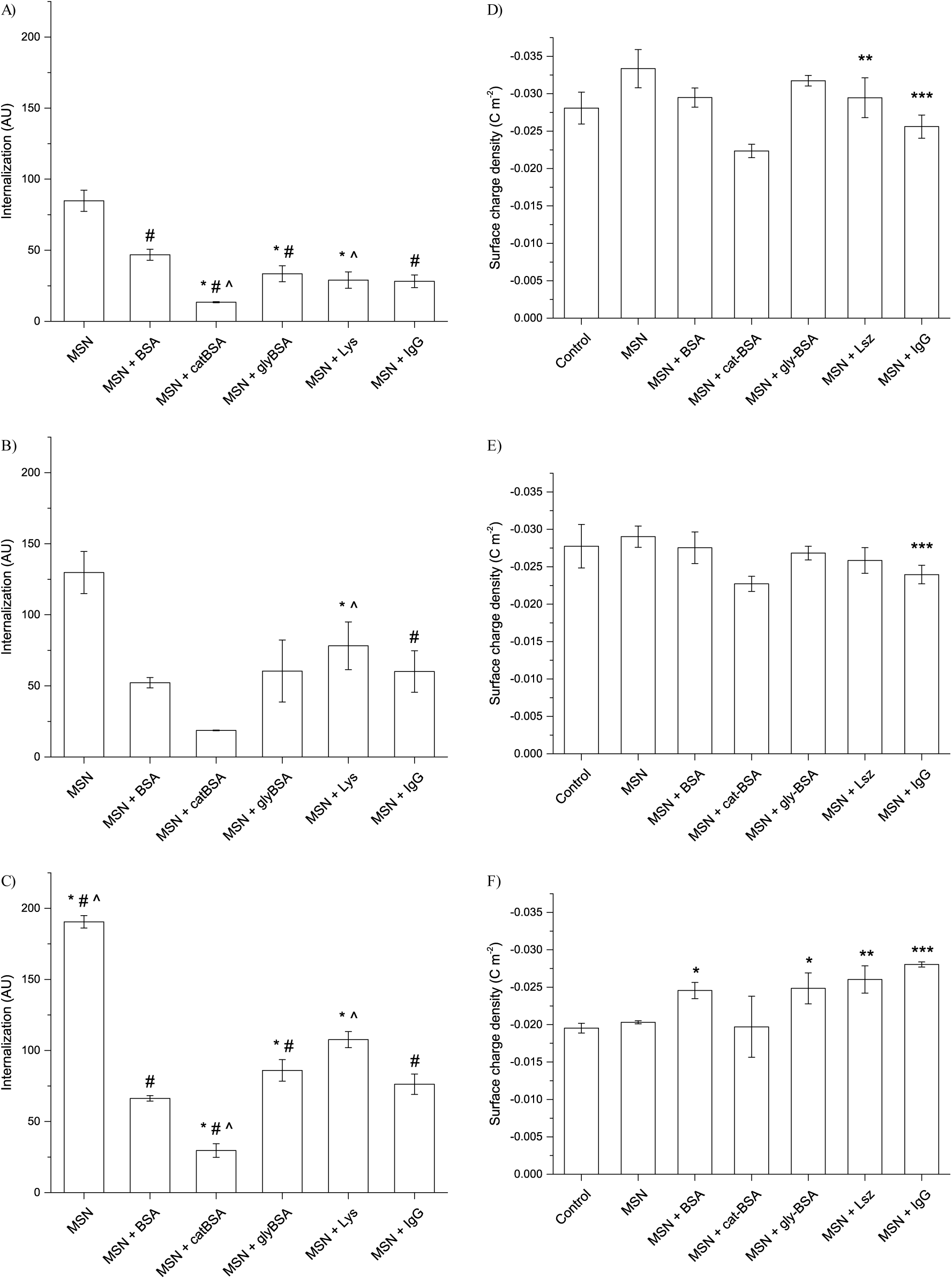
Cellular internalization of protein-coated mesoporous silica nanoparticles (MSN) after 6 hours of exposure by A) NIH/3T3 fibroblasts, B) EAC-E carcinoma cells, and C) RAW264.7 macrophages. *, # and ^ indicate statistically significant difference (p<0.01) with respect to BSA, Lsz and IgG-coated MSN groups respectively. Surface charge density of cell membranes after exposure to protein-coated MSN after 6 hours of exposure. Changes in surface charge density of D) NIH/3T3 fibroblasts, E) EAC-E carcinoma cells, and F) RAW264.7 macrophages. * indicates statistically significant difference with respect to the corresponding control cells (* p<0.05, ** p<0.01, *** p<0.001).

Differential cellular internalization may affect cellular response to protein-coated nanoparticles. Cellular interactions of nanoparticle begin at the cell membrane. Nanoparticle interaction is known to compromise the integrity of cell membranes [49]. However, despite the knowledge that endocytosis results in changes in the composition of the cell membrane [50], there is insufficient understanding of the effect of nanoparticle interaction on the cell membrane composition at sub-toxic conditions. The protein corona, which acts as an intermediary between the nanoparticle surface and the cell membrane, may play an essential role in the modulation of these cell membrane compositional changes. Changes in the composition and functional groups often result in an altered surface charge density of the cell membrane due to differences in the concentrations of charged chemical groups such as amines, phosphates, carboxyls, and sulfates [51]. Therefore, to study the cellular response to nanoparticles uptake, we measured the surface charge density of cells as a surrogate to estimate changes in cell membrane functional groups. For estimating the surface charge densities of the three cell types after exposure to different protein-coated MSN, we measured the electrophoretic mobility of the cell surface under externally applied electric field (Figure 6D-F). The surface charge density was negative for all cell types due to negatively charged phosphate groups present on cell membrane phospholipids [51]. We observed cell type-dependent differences due to the interaction with different protein-coated MSN. For fibroblasts, the surface charge density significantly decreased after interaction with bare MSN, which may be due to a high amount of internalization of this group compared to the protein-coated MSN (Figure 6D). On the other hand, cat-BSA-coated MSN, despite being internalized in meagre amounts, significantly increased the surface charge density of fibroblasts. This increase in surface charge of fibroblast is not an effect of the cationic surface of cat-BSA as Lsz-coated MSN did not result in a similar effect. It may, instead, be an effect of interaction with large-sized agglomerates of cat-BSA-coated MSN which induced significant changes in the cell membrane composition without getting internalized. We also observed an increase in the surface charge density of carcinoma cells exposed to cat-BSA-coated MSN (Figure 6E). However, carcinoma cells were less susceptible to changes in surface charge density compared to fibroblasts after exposure to all other protein-coated MSN.

The effect of interaction with protein-coated MSN on surface charge density was markedly distinct for macrophages compared to the other two types of cells (Figure 6F). We observed significant reductions in surface charge density of macrophages after exposure to all protein-coated MSN except for cat-BSA-coated MSN. Unlike the other cell types, the cell membrane of macrophages was not as affected by cat-BSA-coated MSN agglomerates. Furthermore, IgG-coated MSN induced the most significant reduction in surface charge density for macrophages. We speculate it may be due to selective recognition of the Fc-region of IgG bound to the MSN surface and consequent cellular responses [52].

Overall, these results indicate that protein physicochemical properties affect both nanoparticle-protein and nanoparticle-cell interaction. The surface characteristics of proteins, especially surface charge, had a significant effect on the binding affinity and amount of protein adsorbed on MSN. The secondary structure of protein had a significant effect on structural rearrangements. The colloidal state of the protein-MSN complex formed as a result of nanoparticle-protein interactions significantly influenced the nanoparticle uptake by cells. The uptake was characteristic for different cells and caused changes in the plasma membrane composition as estimated by cell surface charge density. In particular, the response of macrophage cells to nanoparticles is significantly dependent on the properties of the protein corona, which may have implications in the biodistribution of nanoparticles introduced into the bloodstream for biomedical applications. The future studies about nano-bio interface should include serum from individuals with various pathophysiological conditions to develop safe and efficacious nanotherapeutic interventions.

## Supporting information

