## Supplementary material for "Role of Physicochemical Properties of Protein in Modulating the Nanoparticle-Bio interface"

#### Supplementary contents

### 1 Supplementary Figures

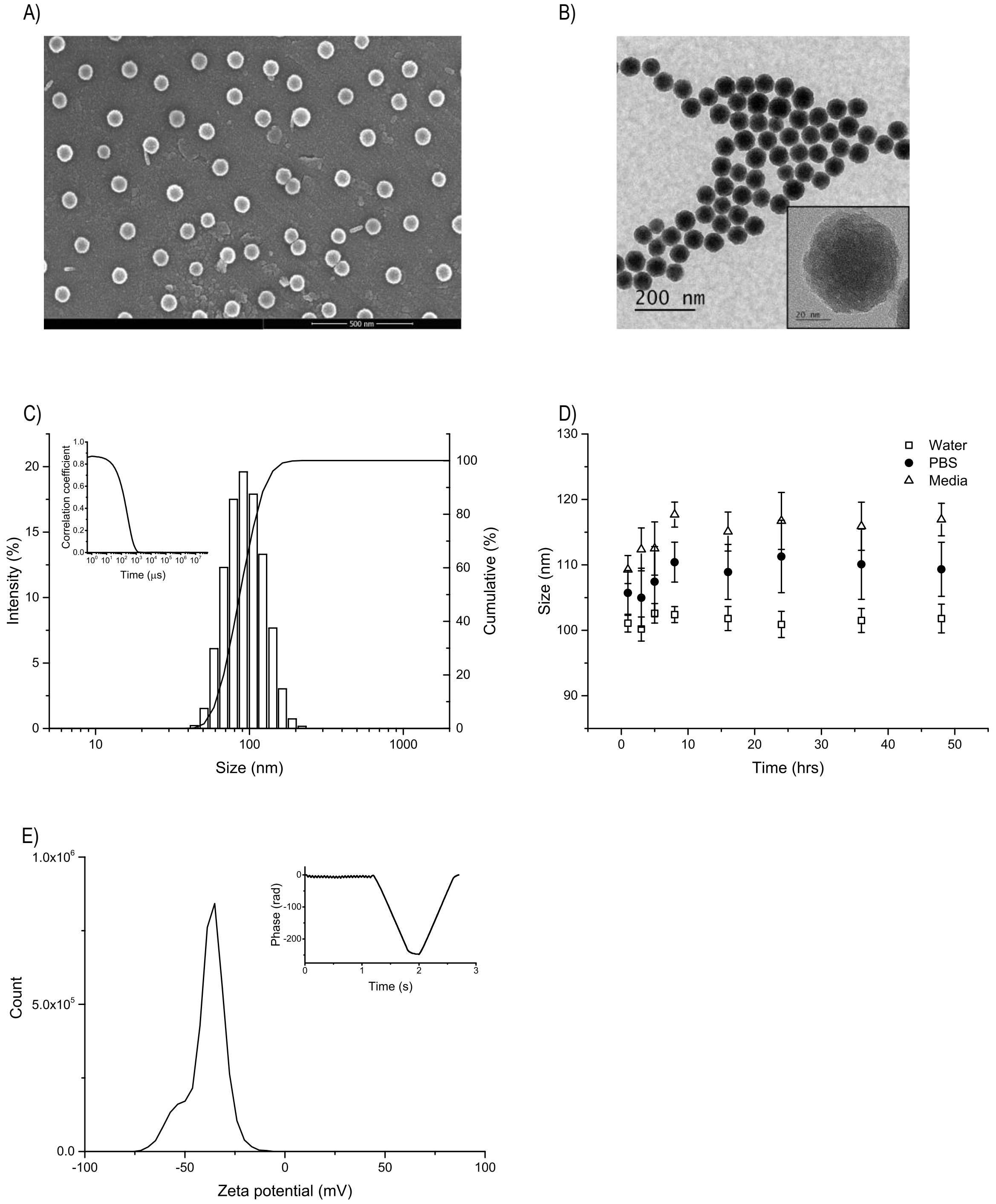

**Figure S1. Characterization of synthesized mesoporous silica nanoparticles (MSN).** A) Scanning electron micrograph (SEM) of MSN (scale bar – 500 nm). B) Transmission electron micrograph (TEM) of MSN (scale bar – 200 nm), with the inset showing a high magnification micrograph of a single nanoparticle (scale bar – 20 nm). C) Hydrodynamic size distribution of MSN measured using dynamic light scattering (DLS), with the inset showing the intensity decay correlogram. D) Hydrodynamic size of MSN suspended in water, phosphate buffered saline (PBS) and media over a period of 48 hours. E) Zeta potential distribution of MSN measured using laser Doppler velocimetry (LDV), with the inset showing the phase plot.

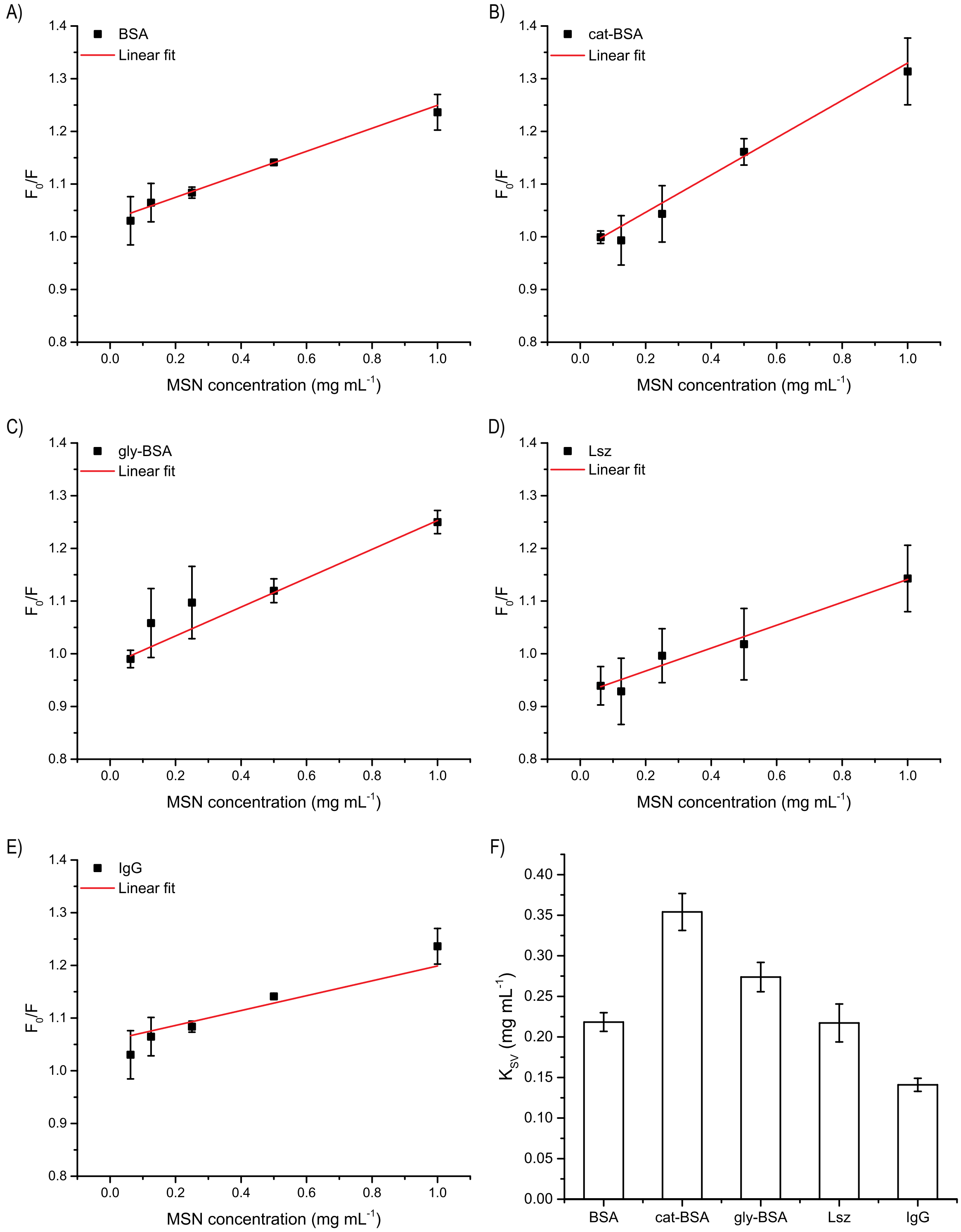

**Figure S2. Fluorescence quenching of proteins due to MSN interaction.** A-E) Ratio of fluorescence intensities of native and MSN-bound BSA (A), cat-BSA (B), gly-BSA (C), Lsz (D) and IgG (E) as a function of MSN concentration. The red lines represent the linear fit of the data. F) Stern-Volmer fluorescence quenching constants of proteins due to MSN interaction obtained from the linear fit of the data.

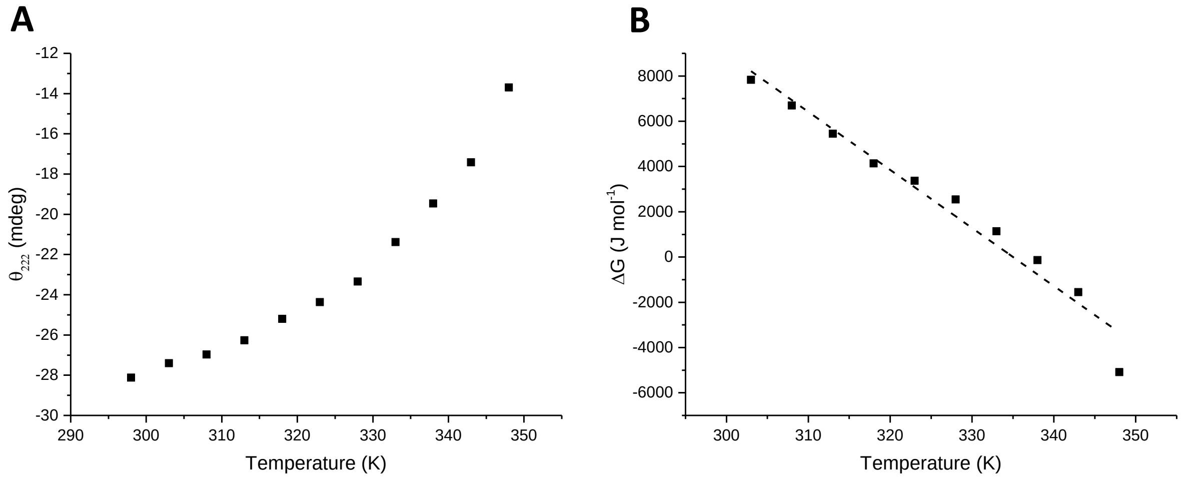

**Figure S3. Temperature-dependent CD spectroscopy of native BSA.** A) Change in ellipticity values at 222 nm ($\theta_{222}$) as a function of temperature. B) Change in energy of unfolding ($\Delta G$) as a function of temperature. The dashed line represents the linear fit of the data.

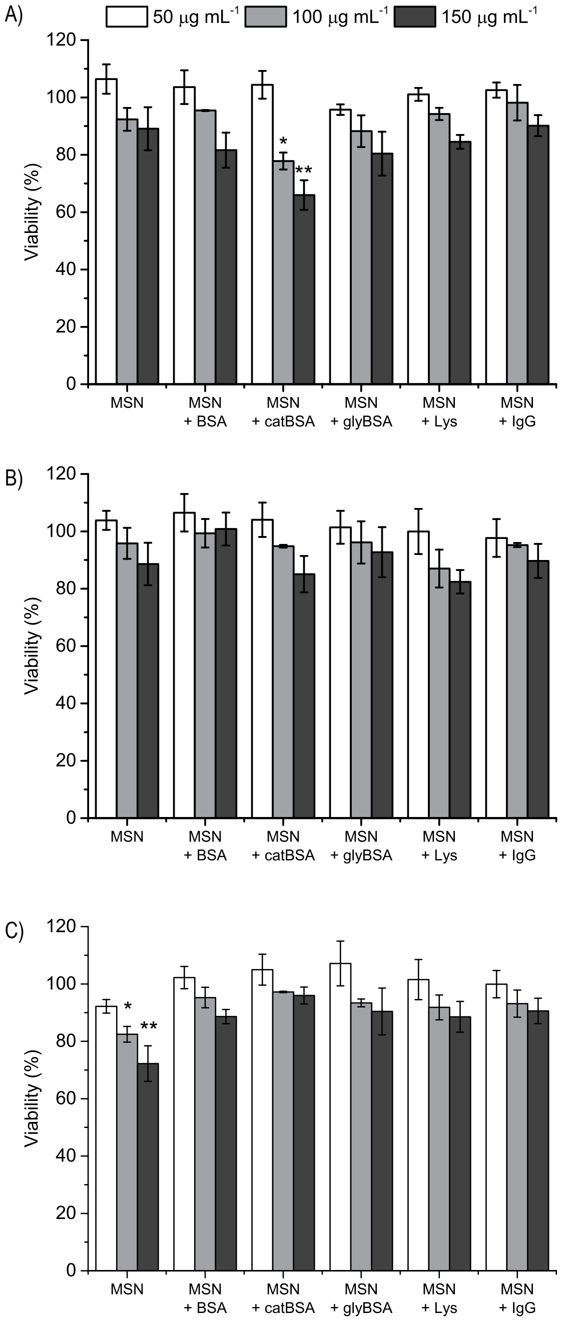

**Figure S4. Cytotoxicity of protein-coated MSN.** Cellular viability of A) NIH/3T3 fibroblasts, B) EAC-E carcinoma cells and C) RAW264.7 macrophages exposed to different concentrations of protein-coated MSN. * indicates statistically significant difference with respect to the corresponding control (untreated) cells (* $p<0.001$, ** $p<0.0001$).

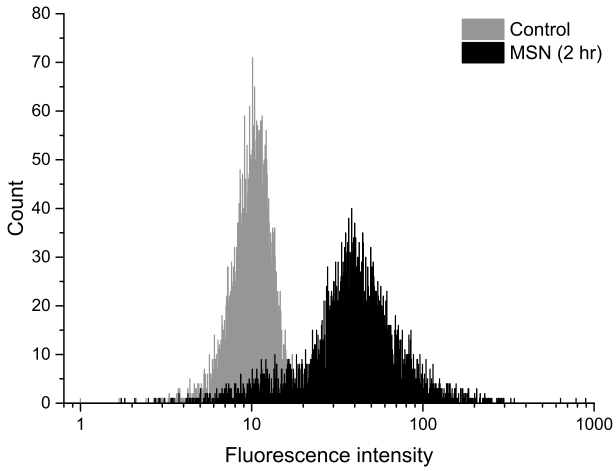

**Figure S5. Flow cytometry histogram of NIH/3T3 cells.** Fluorescence intensity distributions of cells incubated in control media (grey) and in media containing MSN (black) for 2 hrs.

#### 2 Supplementary Tables

**Table S1.** Two-way ANOVA analysis of secondary structure of model proteins (n=3). The values reported are adjusted p-values calculated using Tukey's multiple comparison test. (BSA – Bovine Serum Albumin, cBSA – cationized BSA, gBSA – glycated BSA, Lsz – Lysozyme, and IgG – immunoglobulin G). Values in bold are statistically significant.

| | | **BSA** | **cat-BSA** | **gly-BSA** | **Lsz** | **IgG** |
|---|---|---|---|---|---|---|
| **Helices** | **BSA** | | 0.5915 | 0.5754 | **<0.0001** | **<0.0001** |
| | **cat-BSA** | | | >0.9999 | **<0.0001** | **<0.0001** |
| | **gly-BSA** | | | | **<0.0001** | **<0.0001** |
| | **Lsz** | | | | | **<0.0001** |
| **Sheets** | | **BSA** | **cat-BSA** | **gly-BSA** | **Lsz** | **IgG** |
| | **BSA** | | **0.0373** | 0.8165 | **<0.0001** | **<0.0001** |
| | **cat-BSA** | | | 0.3340 | **<0.0001** | **<0.0001** |
| | **gly-BSA** | | | | **<0.0001** | **<0.0001** |
| | **Lsz** | | | | | **<0.0001** |
| **Turns** | | **BSA** | **cat-BSA** | **gly-BSA** | **Lsz** | **IgG** |
| | **BSA** | | >0.9999 | 0.9855 | **<0.0001** | **<0.0001** |
| | **cat-BSA** | | | 0.9938 | **<0.0001** | **<0.0001** |
| | **gly-BSA** | | | | **<0.0001** | **<0.0001** |
| | **Lsz** | | | | | >0.9999 |
| **Unordered** | | **BSA** | **cat-BSA** | **gly-BSA** | **Lsz** | **IgG** |
| | **BSA** | | 0.4954 | >0.9999 | **0.0121** | **<0.0001** |
| | **cat-BSA** | | | 0.5432 | **0.0001** | **<0.0001** |
| | **gly-BSA** | | | | **0.0099** | **<0.0001** |
| | **Lsz** | | | | | 0.0767 |

**Table S2.** One-way ANOVA analysis of protein adsorption (n=3). P-values of Tukey's multiple comparison tests are reported for pair-wise comparison of data. Values in bold are statistically significant.

| | | BSA | cat-BSA | gly-BSA | Lsz |
|---|---|---|---|---|---|
| **$A_{max}$** | BSA | | | | |
| | cat-BSA | **<0.0001** | | | |
| | gly-BSA | **0.0445** | **0.0001** | | |
| | Lsz | **0.0001** | **0.0495** | **0.0083** | |
| | IgG | **<0.0001** | **0.0010** | **<0.0001** | **<0.0001** |
| **$k_D$** | BSA | | | | |
| | cat-BSA | **0.0259** | | | |
| | gly-BSA | 0.7186 | **0.0039** | | |
| | Lsz | **<0.0001** | **0.0037** | **<0.0001** | |
| | IgG | **<0.0001** | **<0.0001** | **0.0002** | **<0.0001** |

**Table S3.** One-way ANOVA analysis of Stern-Volmer fluorescence quenching constants of proteins after interaction with MSN (n=3). P-values of Tukey's multiple comparison tests are reported for pair-wise comparison of the data. Values in bold are statistically significant.

| | **BSA** | **cat-BSA** | **gly-BSA** | **Lsz** |
|---|---|---|---|---|
| **BSA** | | | | |
| **cat-BSA** | **<0.0001** | | | |
| **gly-BSA** | **0.0223** | **0.0018** | | |
| **Lsz** | >0.9999 | **<0.0001** | **0.0198** | |
| **IgG** | **0.0024** | **<0.0001** | **<0.0001** | **0.0027** |

**Table S4.** Hydrodynamic diameter and zeta potential of PC-MSN. # indicates agglomeration of the PC-MSN complex.

| **Group** | **Hydrodynamic diameter** (nm) | **Zeta potential** (mV) |
|---|---|---|
| MSN (bare) | 90.2 ± 0.7 | -42.2 ± 1.3 |
| MSN + BSA | 100.6 ± 1.7 | -32.9 ± 5.1 |
| MSN + cat-BSA | 2160.3 ± 236.0 # | 4.2 ± 1.4 |
| MSN + gly-BSA | 108.3 ± 0.7 | -28.1 ± 7.7 |
| MSN + Lsz | 139.1 ± 4.5 | 20.6 ± 0.7 |
| MSN + IgG | 1237.3 ± 163.8 # | 1.5 ± 0.7 |

**Table S5.** Two-way ANOVA for ellipticity of proteins in Figure 3 (contributions of temperature and MSN interaction)

| | Parameters | Helices | Sheets | Turns | Unordered |
|---|---|---|---|---|---|
| **BSA** | **Interaction** | 0.404% (0.8983) | 3.173% **(0.0088)** | 7.811% **(0.0243)** | 8.874% **(0.0445)** |
| | **Temp** | 95.47% **(<0.0001)** | 91.11% **(<0.0001)** | 76.23% **(<0.0001)** | 69.19% **(<0.0001)** |
| | **MSN** | 0.5466% **(0.0093)** | 0.5731% **(0.0251)** | 1.053% (0.0717) | 2.966% **(0.0086)** |
| **cBSA** | **Interaction** | 0.8671% (>0.9999) | 6.67% (0.3361) | 16.46% **(<0.0001)** | 13.22% **(0.007)** |
| | **Temp** | 31.54% **(0.0173)** | 44.73% **(<0.0001)** | 71.33% **(<0.0001)** | 41% **(<0.0001)** |
| | **MSN** | 10.76% **(0.0041)** | 23.61% **(<0.0001)** | 0.6601% (0.1042) | 25.07% **(<0.0001)** |
| **gBSA** | **Interaction** | 0.9994% (0.7966) | 0.9521% (0.7219) | 5.182% (0.3364) | 15.00% (0.2772) |
| | **Temp** | 91.99% **(<0.0001)** | 92.56% **(<0.0001)** | 74.76% **(<0.0001)** | 32.93% (**0.0074**) |
| | **MSN** | 0.05624% (0.5362) | 0.6449% **(0.0257)** | 0.6285% (0.2187) | 0.02486% (0.8803) |
| **Lysozyme** | **Interaction** | 18.57% (0.242) | 11.54% (0.1002) | 18.39% (0.214) | 18.05% (0.1711) |
| | **Temp** | 5.641% (0.9486) | 53.52% **(<0.0001)** | 23.02% (0.0962) | 27.54% **(0.0262)** |
| | **MSN** | 14.48% **(0.0015)** | 5.44% **(0.0046)** | 0.3455% (0.596) | 1.105% (0.3236) |
| **IgG** | **Interaction** | 21.39% **(0.0091)** | 12.13% (0.2213) | 16.62% (0.3517) | 8.332% (0.9219) |
| | **Temp** | 43.72% **(<0.0001)** | 46.46% **(<0.0001)** | 19.04% (0.2491) | 9.642% (0.8764) |
| | **MSN** | 0.03215% (0.8342) | 2.574% (0.0808) | 0.8083% (0.4383) | 1.961% (0.2836) |

**Table S6.** Three-way ANOVA for internalization of PC-MSN (contributions of cell type, protein type and exposure time)

| Source | Degrees of freedom | F value | P value | % of total variation |
|---|---|---|---|---|
| Cell type | 2 | 171.354 | **<0.001** | 25.31 |
| Time | 3 | 8.992 | **<0.001** | 1.99 |
| Protein corona | 5 | 63.513 | **<0.001** | 23.45 |
| Cell type × Time | 6 | 2.700 | **0.016** | 1.20 |
| Cell type × Protein corona | 10 | 39.921 | **<0.001** | 29.48 |
| Time × Protein corona | 15 | 1.551 | 0.095 | 1.72 |
| Cell type × Time × Protein corona | 30 | 2.801 | **<0.001** | 6.21 |

### 3 Supplementary Methods

#### 3.1 Synthesis and characterization of mesoporous silica nanoparticles

##### 3.1.1 Synthesis of mesoporous silica nanoparticles

Mesoporous silica nanoparticles (MSN) were synthesized using a water-in-oil micro-emulsion technique as described previously [1]. Briefly, the aqueous phase (800 µL of ultra-pure type I water) was dispersed in an organic phase of cyclohexane (15.4 mL), Triton X-100 (3.54 g) (surfactant), and hexanol (3.2 mL) (co-surfactant). Tetraethyl orthosilicate (TEOS) (400 µL) was then added as a silane precursor, along with aqueous ammonia (400 µL) which acts as a catalyst for the hydrolysis of TEOS. Subsequently, 3-(trihydroxysilyl) propylmethylphosphonate (THPMP) (30 µL) was added to modify the nanoparticles with methyl phosphonate groups to obtain an inert nanoparticle surface to prevent agglomeration. The emulsion was stirred in a round-bottom flask for 24 hrs at room temperature, following which the emulsion was broken by adding absolute ethanol. The nanoparticles were separated using centrifugation at 14000 RCF for 30 mins, and were then washed thrice in absolute ethanol and thrice in ultra-pure (type I) water.

Fluorescently-tagged MSN were synthesized using a conjugate of tetramethylrhodamine-isothiocyanate (TRITC) and (3-aminopropyl)triethoxysilane (APTES). TRITC-APTES conjugate (50 µL) was added to the micro-emulsion before the addition of silane precursor (TEOS) in the nanoparticle synthesis procedure as stated above.

##### 3.1.2 Quantification of nanoparticle concentration

The concentration of nanoparticle suspension was quantified by measuring the dry weight of a unit volume of MSN suspension. The nanoparticle suspension was centrifuged at 14000 RCF for 30 mins to obtain a nanoparticle pellet, following which the supernatant was discarded and the pellet was dried under vacuum at 70°C for 7-8 hrs.

##### 3.1.3 Morphology of nanoparticles

The morphology and the size of the nanoparticles in a dry state were studied using scanning electron microscopy. MSN were suspended in absolute ethanol at a concentration of 100 $\mu g \cdot mL^{-1}$. A 10 µL of ethanol suspension of MSN was dried on a sheet of clean aluminium foil and then placed on a copper stub using carbon tape. The sample was kept under vacuum overnight for drying. Gold sputter coating was then performed on the sample, followed by microscopy using a Nova NanoSEM 450 (FEI, Oregon, USA) at 100,000X magnification.

Morphology and size were also characterized using transmission electron microscopy. Samples were prepared by placing a 10 µL of aqueous suspension of MSN on a Formvar-coated 300 mesh copper TEM grid (Ted Pella Inc., California, USA) and drying for 48 hrs, following by microscopy using a Tecnai G2 12 Twin TEM (FEI, Oregon, USA).

Image analysis was performed to estimate the average diameter of MSN using ImageJ software [2].

##### 3.1.4 Hydrodynamic size of nanoparticles

The hydrodynamic diameter of nanoparticles in suspension was determined by the dynamic light scattering (DLS) technique using a Zetasizer ZS90 (Malvern Instruments, Malvern, UK). The mobility of suspended nanoparticles was determined by measuring fluctuations in intensity of scattered light, from which the size distribution of nanoparticles was calculated using an auto-correlation function.

##### 3.1.5 Surface charge of nanoparticles

The surface charge of nanoparticles was determined by measuring the zeta potential of nanoparticle surface in aqueous suspension using laser Doppler velocimetry (LDV) measurements using a Zetasizer ZS90.

##### 3.1.6 Hydrodynamic size and surface charge of protein-coated nanoparticles

MSN suspension (1.0 mg·mL$^{-1}$) was mixed with equal volume of protein solution (0.25 mg mL$^{-1}$) in Eppendorf tubes and incubated at 37°C for 24 hrs. DLS and LDV measurements were performed using a Zetasizer ZS90 to determine the hydrodynamic diameter and zeta potential of the protein-coated nanoparticles.

#### 3.2 Fluorescence quenching assay

Autofluorescence of proteins in free state and in the presence of increasing MSN concentrations was measured using synchronous scanning fluorescence spectroscopy [3]. Protein samples (50 µg·mL$^{-1}$) were incubated in presence or absence of MSN (3.125 to 50 µg·mL$^{-1}$) for 24 hrs at 37°C, following which 100 µL of the sample was taken in a non-binding, black opaque 96-well plate (Corning Inc., New York, USA) and fluorescence intensity was detected using a spectrophotometer (Synergy H4, BioTek Instruments, Vermont, USA). The fluorescence of tryptophan residues in the proteins was measured by scanning the excitation wavelength from 250 to 340 nm, and the emission wavelength from 310 to 400 nm. A fixed wavelength difference of 60 nm between excitation and emission was maintained in order to resolve tryptophan. Nanoparticles were considered to be acting as quenching molecules, and the data was fitted to the Stern-Volmer model of fluorescence quenching using equation 2.

$$\frac{F_0}{F} = 1 + K_{SV}[MSN] \tag{2}$$

Here,

$F_0$ is fluorescence intensity of free protein (AU),

$F$ is fluorescence intensity of MSN-bound protein (AU),

$[MSN]$ is the concentration of nanoparticles in suspension (mg·mL$^{-1}$), and

$K_{SV}$ is the Stern-Volmer fluorescence quenching constant (mL·mg$^{-1}$).

#### 3.3 Cytotoxicity of nanoparticles

Cytotoxicity of nanoparticles was studied using the Resazurin assay. Cells were seeded on 96-well plates. The media was then aspirated, cells were then washed with 1X PBS, and protein-coated MSN suspension in serum-free media was added. After 24 hrs of exposure, the nanoparticle suspension was aspirated and cells were washed with 1X PBS three times to remove any remaining nanoparticles. Resazurin dye 20 $\mu g \cdot mL^{-1}$ in complete media was then added to the cells and incubated at 37°C for 5 hrs. Resazurin is reduced by viable cells to resorufin whose fluorescence is measured at excitation wavelength 540 nm and emission wavelength 600 nm, with a 20 nm bandpass.
